# High-speed motility originates from cooperatively pushing and pulling flagella bundles in bilophotrichous bacteria

**DOI:** 10.1101/629121

**Authors:** Klaas Bente, Sarah Mohammadinejad, Mohammad A. Charsooghi, Felix Bachmann, Agnese Codutti, Christopher T. Lefèvre, Stefan Klumpp, Damien Faivre

## Abstract

Bacteria propel and change direction by rotating long, helical filaments, called flagella. The number of flagella, their arrangement on the cell body and their sense of rotation hypothetically determine the locomotion characteristics of a species. The movement of the most rapid microorganisms has in particular remained unexplored because of additional experimental limitations. We show that magnetotactic cocci with two flagella bundles on one pole swim faster than 500 µm⋅s^−1^ along a double helical path, making them one of the fastest natural microswimmers. We additionally reveal that the cells reorient in less than 5 ms, an order of magnitude faster than reported so far for any other bacteria. Using hydrodynamic modeling, we demonstrate that a mode where a pushing and a pulling bundle cooperate is the only possibility to enable both helical tracks and fast reorientations. The advantage of sheathed flagella bundles is the high rigidity, making high swimming speeds possible.

## Introduction

The understanding of microswimmer motility has implications ranging from the comprehension of phytoplankton migration to the autonomously acting microbots in medical scenarios [1, 2]. The most present microswimmers in our daily lives are bacteria, most of which use flagella for locomotion. Well-studied examples of swimming microorganisms include the peritrichous (several flagella all over the body surface) *Escherichia coli* with an occasionally distorted hydrodynamic flagella bundling [3] and the monotrichous (one polar flagellum) *Vibrio alginolyticus*, which are pushed or pulled by a flagellum and exploit a mechanical buckling instability to change direction [4, 5]. The swimming speeds of so far studied cells are in the range of several 10 µm s^−1^ and their reorientation events occur on the time scale of 50-100 ms [5, 6].

*Magnetococcus marinus* (MC-1) is a magnetotactic, spherical bacterium that is capable of swimming extremely fast [7–10]. MC-1 as well as the closely related strain MO-1 [11] are equipped with two bundles of flagella on one hemisphere (bilophotrichous cells). The bacterium also features a magnetosome chain, which imparts the cell with a magnetic moment (‘magnetotactic’ cell). They are assumed to swim with the cell body in front of both flagella, which synchronously push the cell forward [7]. This assumption leads to helical motion in the presence of a strong magnetic field, which exerts a torque on the cell’s magnetic moment, as seen in hydrodynamic simulations [12]. In the absence of a magnetic field, this model predicts rather straight trajectories.

Our observations disagree with the above-mentioned model, indicating that an understanding of the physics of their swimming is still missing, even though proof of concept biomedical applications of these bacteria have already emerged [2]. Here we show that MC-1 not only reach speeds of over 400 µm s^−1^ but that this speed is recorded along an unexplored double helical path. In addition, this rapid movement is also complemented by an extremely fast reorientation ability (less than 5 ms). We correlate the flagella architecture and the swimming mechanism by hydrodynamic simulations and show that only a striking cooperative movement where one flagella bundle pushes while the other pulls the cell explains these motility characteristics.

## Results

We observed MC-1 cells in a physicochemically controlled environment that resembled the cell’s natural habitat [13] (Fig. 1A). The cells were introduced into a flat, rectangular microcapillary tube with a height of 200 µm, where they accumulated in a band near their preferred oxygen conditions. The cells were imaged near this band [14, 15] and at central height to avoid surface interactions. The capillary was placed at the center of three orthogonal Helmholtz coils [16], which were used to cancel the Earth’s magnetic field with a precision of 0.2 µT after the band had formed. Hence, the cells’ motion could be observed in the absence of magnetic torques. Tracking was performed in 3D at 400 frames s^−1^ (fps) (Fig. 1B). A high-throughput tracking method was used [17] (see Methods) for the reconstruction of the tracks (Fig. 1C and D).

**Figure 1.**
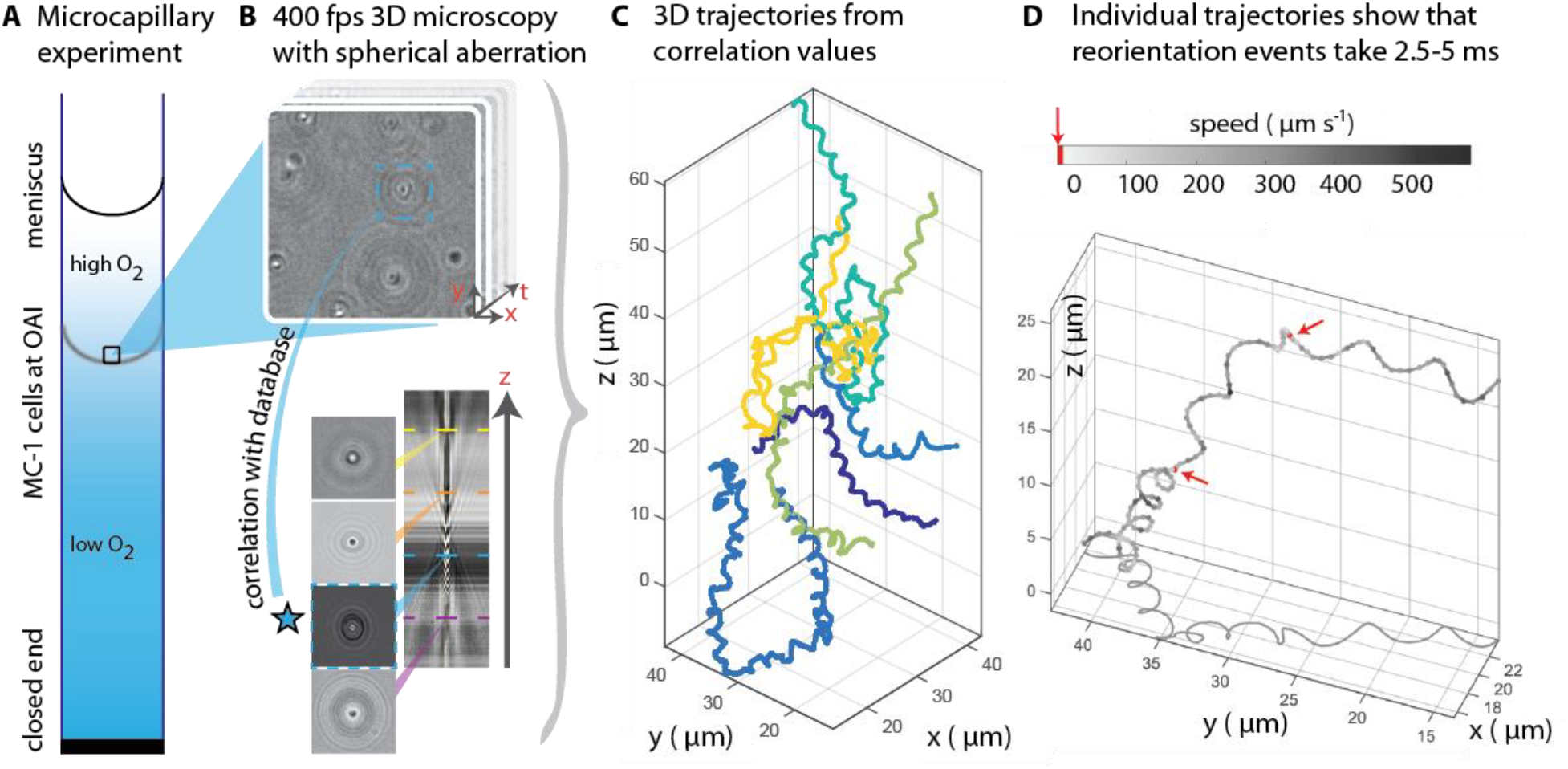
3D tracking in physiochemically controlled conditions. **A** MC-1 cells were transitioned into agar-free medium, the solution was degassed to remove the oxygen and inserted into a microcapillary. Due to one open and one closed end, an oxygen gradient formed and the cells accumulated near their preferred microoxic conditions (the oxic-anoxic-interface, OAI). **B** The cells were observed near the band using 400 fps phase contrast video microscopy with a spherical aberration, which causes interference patterns around the spherical swimmers. These patterns can be correlated with patterns from silica beads of known height relative to the focal plane. **C** A tracking algorithm enables high-throughput 3D tracking of the microswimmers [17]. Colors indicate different cells. **D** Individual tracks were analyzed and a clockwise helical travel path with a radius close to the cell diameter was found as well as instantaneous traveling speeds between 100 µm s^−1^ and 500 µm s^−1^. Tracks can be interrupted by rapid reorientation events that last only 2.5-5 ms. The helix parameters like pitch and period time do not change before and after an event, but apparently do so in the projected 2D tracks (see projected shadow in **D**).

Our first set of observations revealed that the cells traveled on helical paths (with a pitch of 5.3 µm ± 1.3 µm, a diameter of 1.7 µm ± 0.2 µm, and a period of 46 ms ± 32 ms, n = 65) in the absence of magnetic torques. In addition, abrupt changes of the direction of the helical axis were observed (around 90°, see Fig.S3). These directional changes occur within 2.5 to 5 ms, at least one order of magnitude faster than any previously analyzed reorientation event [5, 6]. Directional changes did not occur via a continuous modulation of the ratio of radius and pitch [18], as it has been observed as a part of the chemotaxis of sperm [19]. Rather, the helix parameters were the same before and after such an event. 3D tracking is essential to obtain this conclusion, as projected 2D tracks exhibit apparent changes in the helix parameters (see projected shadow of the track in Figure 1D and Figure S4).

The cells were further examined at 1640 fps in high-intensity dark-field video microscopy to visualize the cell body and the flagella movement in detail (Fig. 2, methods). An exemplary track of the cell body movement is shown in Fig. 2A together with the tracked velocity. At such frame rates, a more complex movement pattern becomes apparent, which was not detectable during the 3D tracking at 400 fps. The cell track can be represented by a superposition of two helices, a small helix on a large helix (Figure 2B), resulting in a position over time 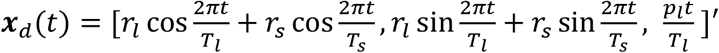. The large helical track features a pitch *p*_*l*_ of ~4 µm, a radius *r*_*l*_ of ~1 µm and a period *T*_*l*_ of 72 ms. The small helix features a pitch of ~0.66 µm, a radius *r*_*s*_ of ~0.125 µm and a period *T*_*s*_ of 14.4 ms. The integer parameter ratio between the period times of the two helices result in a *T*_*l*_–periodic track, indicating a *T*_*l*_–periodic movement pattern of the flagella.

**Figure 2.**
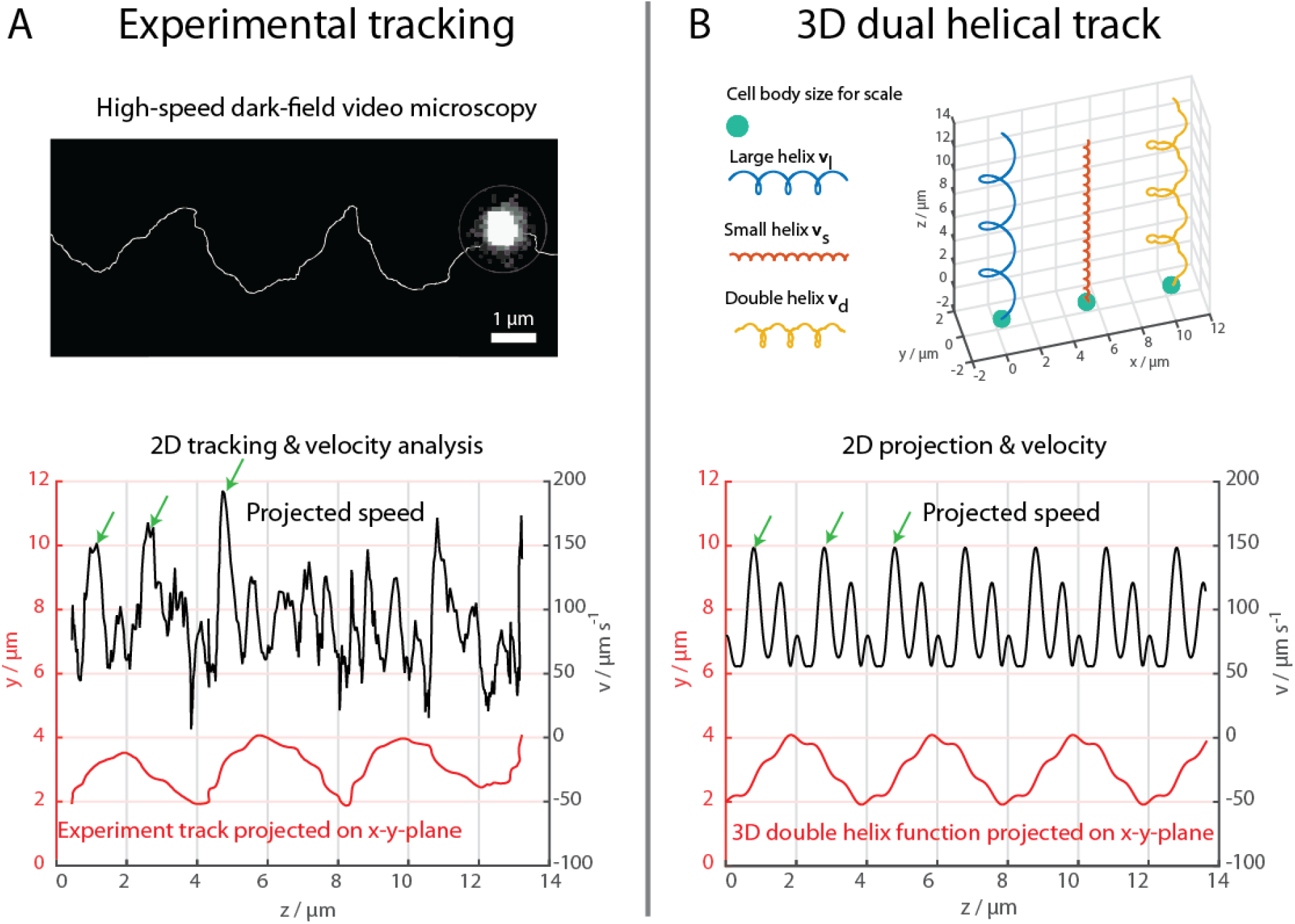
High-speed dark-field video microscopy at 1640 fps reveals a dual helical travel path of a MC-1 cell during free swimming. In A, the tracked path is displayed after smoothing by a 5 point moving average filter and then plotted together with the cell velocity. Green arrows indicate velocity maxima. In B, it is shown that the projected swimming path and projected velocity can be described by a projection of a large 3D double helix 𝑣_d_(*t*). The fact that the ratios between pitch and period time of the large and small helix are integer hints towards a ***T***_***l***_-periodic flagella bundle movement pattern that causes the cell’s swimming behavior.

The flagella movements were also observed in a high-intensity dark-field video microscopy experiments (Fig. 3, and videos S1 & S2). At 1424 fps, short fibers next to the cell body and bright spots on the cell surface could be observed, which we identified as a part of a flagellum close to the cell surface [5]. The cell’s trajectory could be tracked together with the flagella positions on the cell surface over 85 ms, which corresponds to 1.6 periods on the large helical trajectory of the cell. The observed movement pattern is more complex than previously assumed [2, 14, 20, 21]. The two flagella positions move rapidly over the cell body surface. Crucially, one flagellar position is often seen in front of the cell (relative to its swimming direction), contrary to the model of two pushing flagellar bundles, which remain behind the cell body.

**Figure 3.**
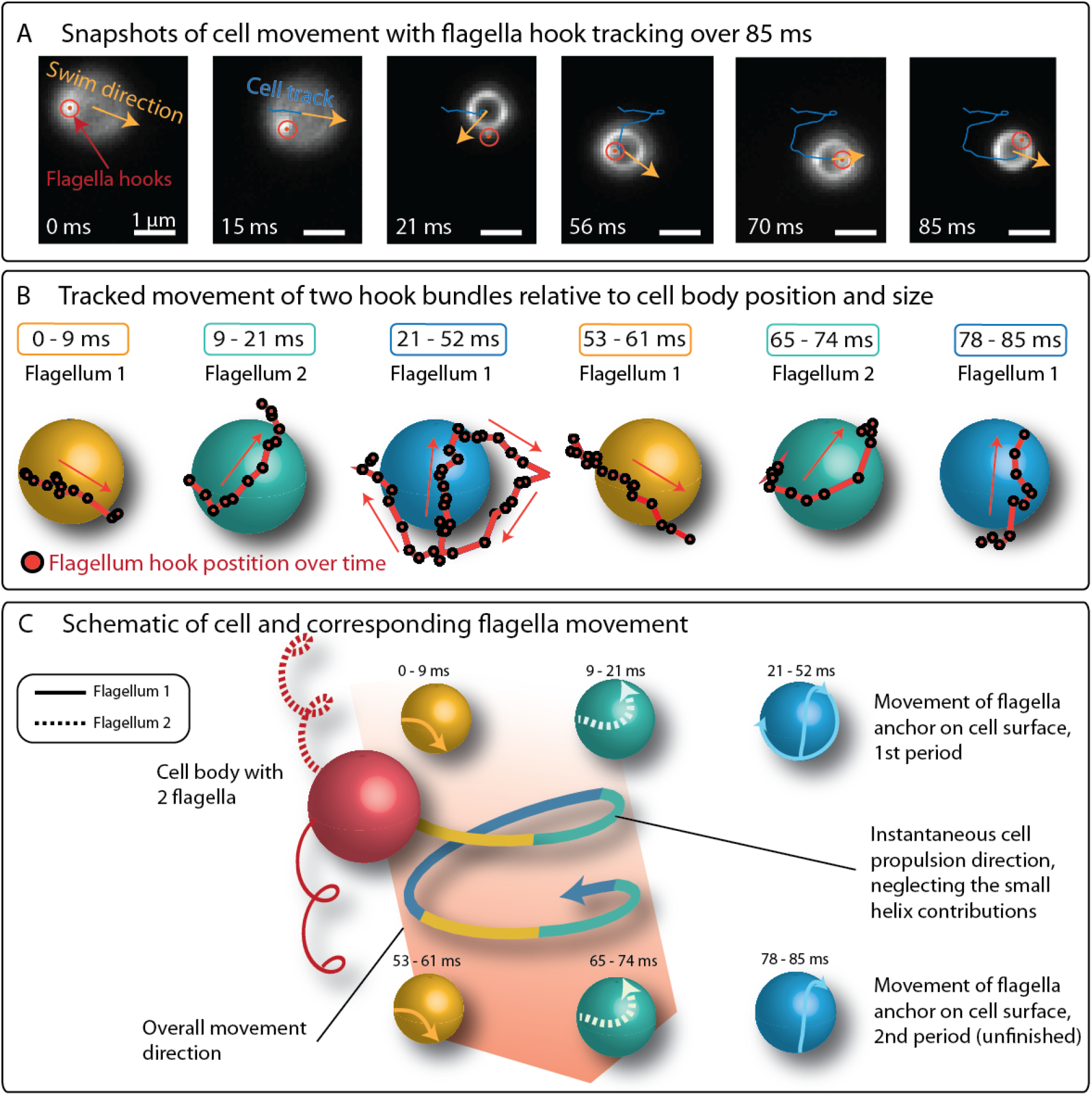
The positions of flagella bundles on the cell surface could be tracked over 85 ms (1.6 large helix period times) along with the cell’s swimming trajectory in high-intensity high-speed dark-field microscopy (from video SI2). **A** While the cell swims from top left to bottom right in the field of view, bright spots can be tracked on the cell surface and directly next to the cell body. The cell’s helical traveling path has a period time of 52 ms. During the cell’s progression, these spots appear to rotate around the spherical cell body with the same periodicity as the cell’s helical path. Due to their apparent distance, their movement pattern, their size and since they can be located above the cell surface in some frames, these spots were identified as parts of the two flagella bundles near the cell body. **B** The positions of both spots were tracked over time, when visible and are depicted relative to a spherical cell body. **C** The color-coding describes the time periods on the large helical track where the flagella were visible. The schematic of the flagella movement pattern appears to be periodic with a period time equal to that of the large helical cell body track.

To test the mechanisms of propulsion and of the rapid reorientations, we turned to simulations of swimming. We performed Stokesian dynamics simulations [22] for a spherical cell body (1.3 µm in diameter) with two discretized helical filaments (4 µm long and 50 nm thick) representing the flagellar bundles (see Methods for details). Since TEM images do not allow for a precise determination of the opening angle between the two flagella (with respect to the body center, see for example Fig. S1), we used a wide range of opening angles (30°-120°) in our simulations. The effect of the flagellar motors was included as a torque at the base of the helices, which rotates the filament, and a counter-torque, which rotates the cell body. We considered the two bundles to rotate independently either counter-clockwise (CCW) or clockwise (CW). The parameters for bending rigidity, torsion rigidity and torque of the flagella were adjusted to reproduce the observed movement characteristics. A motor torque of 12 pN µm, about 3.5 times the motor torque of an *E. coli* cell, had to be chosen together with high isotropic bending and twisting rigidities of 7 pN μm^2^, about 2 times larger than for single flagella [22]. We assume that this increase is attributed to the structure of the flagella bundle. Only this assumption allowed for high swimming velocities in the simulations, indicating that the function of the flagella bundle is to combine high torques with high rigidities.

We simulated the different swimming scenarios arising from combinations of CW and CCW rotation of the two bundles: both flagella pushing the cell (CCW&CCW), both flagella pulling (CW&CW) and one flagellum pushing, one pulling (CCW&CW). The fact that flagella can pull cells is well established [5, 23] and our simulations show that only the CCW&CW model results in the double-helical tracks experimentally observed (Figure 4A & 4B, Figure S5 and Supplementary videos S03 & S04). Tracking the simulated cells results in time traces of the velocity and the projected position that are strikingly similar to the experimental ones (for a quantitative comparison, see Fig. 4 and Fig. 2A). We tested the dependence of the trajectories on model parameters, in particular the opening angle (Supp. Table S1). Double helical trajectories are observed for flagellar opening angles in the whole range we tested, with diameter and pitch of the order of the cell’s body size, in agreement with the experiments.

**Figure 4.**
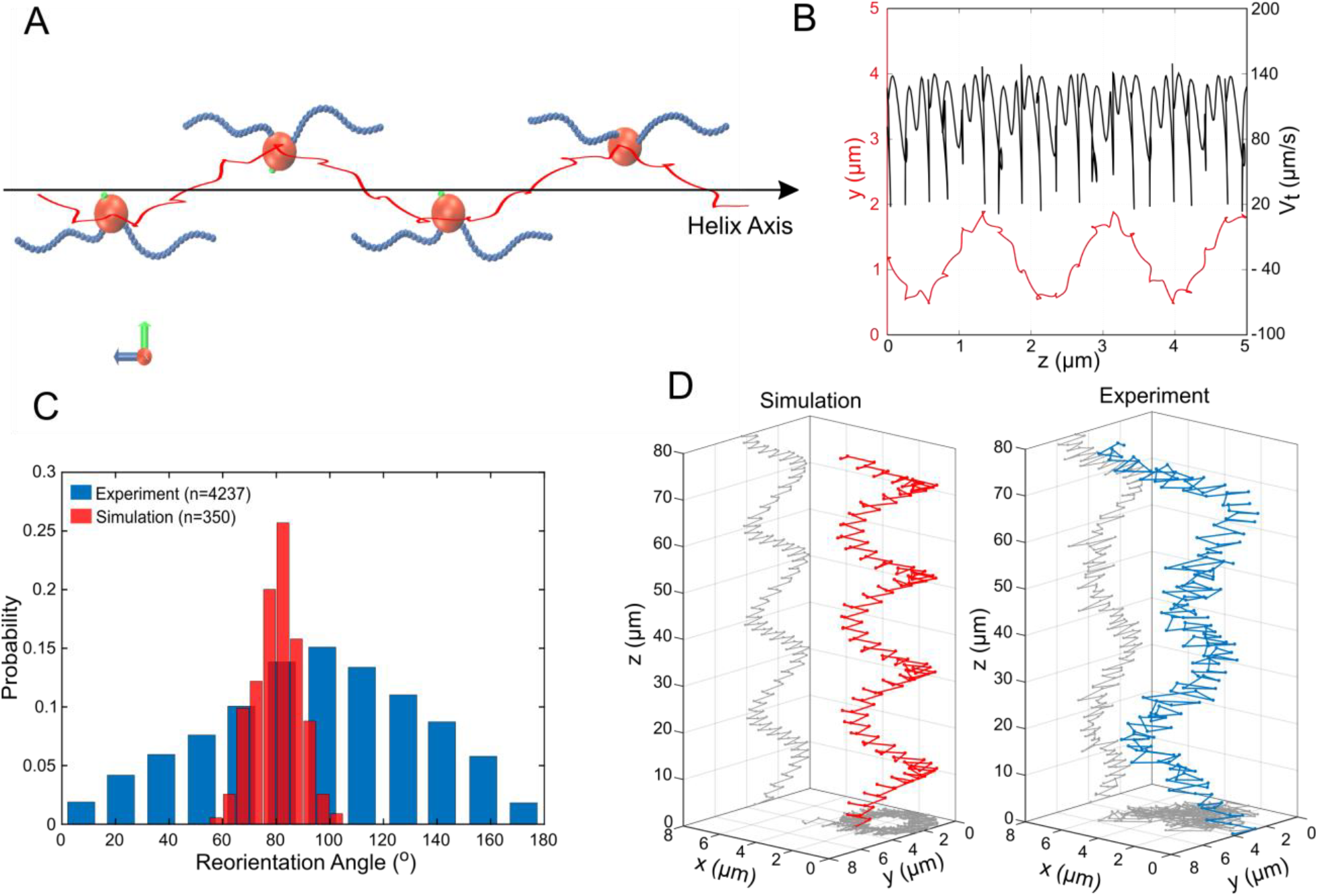
Simulations of MC-1 swimming dynamics with one pushing and one pulling flagellar bundle. The principle cell and flagella arrangement is shown in **A** at four distinct time points where the cell body diverges strongly from the helix axis. **B** shows the projected track of a simulated MC-1 with the flagellar opening angle 60° together with the projected speed. The results are comparable to measured data from Fig. 2A. **C** Histogram of turning angles for the reorientation events seen in experiments and in simulations, where reorientation results from periods of synchronous rotation. **D** Validation of the cooperative pushing and pulling model in the presence of a strong magnetic field (3 mT). A third hyper-helix is observed in both experiment and simulation.

We also tracked the position of the flagella positions on the cell surface in the simulations (by tracking the first beads of each discretized flagellum). Both flagella are seen to rotate around the cell body and one period of the rotation coincides with one period of the large helix in the cell’s trajectory. This agrees qualitatively with the flagella tracking in our experiments and shows that a rotation of the flagella around the cell body causes the regular pattern of the cell’s movement.

The observation that coordination of flagella rotating clockwise and counterclockwise underlies the observed pattern of motion suggests that the rapid reorientation events are due to transiently synchronous rotations of the two flagella. We tested this by changing the sense of rotation from CCW&CW to CCW&CCW for 4 ms in the simulations (outtake in video S05). This procedure indeed resulted in rapid reorientation with an average change in direction by 80° (Figure 4C), in agreement with the 94° change seen in the experiments. The standard deviation of the directional change is small in our simulations (8°), where only the runtime was varied, compared to the experimental value of 39°. The mismatch likely arise from biological diversity in flagella lengths and opening angles, but also due to shifts in local physiochemical conditions, which can e.g. influence the motor torque [5].

To further validate our simulations, we predicted the cell’s swimming behavior at high magnetic fields using our simulation without changing further parameters. The direction of the cell’s magnetic moment was assumed to be perpendicular to the bisector of the two flagella axes (further scenarios can be found in Fig. S5). The simulations showed an additional, large hyper-helical movement pattern (with diameter *D*_sim_ ≃ 3.9 µm and pitch *P*_sim_ ≃ 19.1 µm). The same pattern could thereafter be found in experimental data with similar parameters (*D*_exp_ ≃ 4.2 µm and *P*_exp_ ≃ 30 µm). Typical MC-1 hyper-helical trajectories from simulations and experiments are shown in Figure 4D.

## Discussion

Magnetococci are exceptional swimmers with respect to both their high speed and their reorientation swiftness. In addition, they are likely to make use of a previously unrecognized pattern of motion of their flagella, with one bundle pushing the cell body and the other pulling it. Key to observing this pattern of motion was the 3D tracking of single cells in the absence of magnetic torques to observe the unexpected two-helix trajectories, together with hydrodynamic simulations. The hierarchically organized flagellar bundles provide a high torque and rigidity, necessary to reach record speeds of over 200 body lengths per second, while the unusual type of coordination of the two bundles provides a mechanism for rapid reorientation. Magnetotactic cocci such as MC-1 thrive and generally represent the most abundant MTB in aquatic environments [24]. Their unusual motility is certainly an adaptation that represents a selective advantage that make them competitive in the highly coveted biotopes that are the oxic-anoxic interfaces [25]. Indeed, the observed geometry of this resulting swimming pattern is reminiscent of the situation recently reported for *magnetospirilla* [26], despite the difference in body plan. The *magnetosprilla* have two flagella at opposite cell poles, and a magnetic moment parallel to the flagellar axis, while the flagella of the cocci studied here are attached on one hemisphere of the cell and almost perpendicular to the magnetic moment. Nevertheless, both swim aligned with a magnetic field with one flagellum ahead and one trailing. Record-breaking cells like MC-1 can help to understand physical limits of natural microswimmers and provide design principles for their artificial counterparts.

## Materials and methods

### Cell medium and culturing

MC-1 was cultured similarly to the procedure reported by Bazylinski *et al.* [20]. Artificial sea water (ASW) was used as a base medium, containing 20 g NaCl, 6 g MgCl_2_, 2.4 g Na_2_SO_4_, 0.5 g KCl and 1 g CaCl_2_ per liter H_2_0. To this was added (per liter) the following, in order, prior to autoclaving: 0.05 mL 0.2 % (w/w) aqueous resazurin, 5 mL Wolfe’s mineral solution (ATCC, MD-TMS), 0.3 g NH_4_Cl, 2.4 g HEPES and 1.6 g agar (Kobe I, Carl Roth). The medium was then adjusted to pH 6.3 and autoclaved. After the medium had cooled to about 45 °C, the following solutions were added (per liter), in order, from previously sterile-filtered stock solutions: 0.5 ml vitamin solution (ATCC, MD-VS-10mL), 1.8 mL 0.5 M potassium phosphate buffer, pH 7, 3 mL0.01 M FeCl_2_ and 40 % (w/w) Na thiosulfate. Finally, 0.4 g cysteine was added (per liter), which was made fresh and filter-sterilized indirectly into the medium. The medium (12 mL) was dispensed into sterile Hungate tubes after verifying a pH of 7.0. All cultures were incubated at room temperature (~25 °C) and, after approximately one week, a microaerobic band of MC-1 formed at the oxic–anoxic interface (pink-colorless interface) of the tubes. The cells were harvested in volumes of 1 mL from that region and magnetically transferred to ASW for experiments. The transfer step was necessary to remove agar for swimming experiments and to minimize background scattering in dark-field microscopy.

### Cell morphology analysis

The flagella length was determined with ImageJ from images taken with a Zeiss EM 912 Omega transmission electron microscope using an acceleration voltage of 120 kV. The cells were dried on a carbon film on a regular TEM copper grid and stained with 4 % uranyl acetate for 6 minutes. Due to the staining, the cell walls appeared electron dense and covered the sight on flagella on top or below the cells. Hence, we added the average cell radius to the mean of the flagella length. A mean flagella length of 3.3 µm ± 0.4 µm (n = 27) resulted. The size of the non-dried cells were measured with ImageJ from images taken with a LSM780 (Zeiss; Germany) confocal microscope. The mean size was 1.3 µm ± 0.1 µm (n = 103).

### Microcapillary experiments

1 mL of a freshly harvested sample was degassed using nitrogen for 15 minutes and the sample was introduced into a rectangular micro-capillary (VitroTubes, #3520–050,) by capillary forces. One end of the capillary was sealed with petroleum jelly and the capillary was mounted on a microscope slide that was used to hold the sample on the microscope stage. The oxygen diffusion from the open end caused an oxygen gradient inside the medium, which led together with the oxygen consumption of the cells to the formation of a microaerobic bacteria band. The band formed in the presence of a 50 µT magnetic field towards the sealed end. The tracks were taken after 30 minutes of microcapillary infiltration at 0 µT.

### 3D Tracking experiments

3D swim tracks were recorded at 400 fps in a microcapillary in the vicinity of the microaerobic band (Nikon, S Plan Fluor ELWD, × 40, Ph2, NA 0.6; NA 0.76 condenser lens; Ph2 aperture ring, 635 nm LED illumination). The 3D tracks were reconstructed using the high-throughput phase contrast reference method by Taute *et al.* [17]. A spherical aberration was introduced using a misalignment of the correction collar of the × 40 phase contrast objective to a cover slip thickness correction of 1.2 mm. The aberration caused inference pattern, which can be correlated with the relative height of the microswimmer. A custom made microscope platform, developed by Bennet et al. [16], was used, which features 3 orthogonal Helmholtz coil pairs around the sample position. The setup can generate homogeneous fields at the sample position with arbitrary direction with a precision of 0.2 µT. The Earth’s magnetic field was canceled or an artificial field towards low oxygen conditions was generated during a capillary experiments.

### Dark-field microscopy

Flagella positions were visualized at 1424 fps using high-intensity dark-field microscopy (Nikon 60×, 0.5-1.25 NA CFI P-Fluor oil objective at 0.75 NA; 1.2 NA oil condenser; mercury lamp illumination) and an Andor Zyla 5.5 (10-tap) camera (6.5 μm per pixel). The cells were placed inside a 10 µm deep chamber in ASW. A deeper chamber did not allow for successful dark-field imaging of flagella due to an increase in noise from background scattering. The focal plane was adjusted to the center of the chamber, such that interactions between the observed flagella bundles and the chamber surfaces were avoided. Presumably due to the flagella size and the high rotation speed of the cell and of the flagella, a direct observation of the whole flagella bundles was not successful and only a small section of each flagellum could be visualized. A sub-millisecond exposure time set the requirement for high photon intensity at the sample position. The intensity was increased until the cells melted instantly when swimming into focus. A green filter prevented the melting while still facilitating sufficient brightness. Scrupulous cleanliness at all optical interfaces was obligatory. High-speed cell body tracking could be accomplished at the center of a 200 µm deep chamber. Optimized visualization was achieved at 1640 fps using dark-field microscopy without the critical illumination from flagella tracking (Zeiss 60×, 1.0 NA; 1.2 NA oil condenser; halogen lamp illumination). The dark-field setup did not allow for a cancellation of external magnetic fields. Measurements of the magnetic field at the capillary position yielded B_x_ = −195 µT ± 0.2 µT, B_y_ = 60 µT ± 0.2 µT and B_z_ = 27 µT ± 0.2 µT.

### Cell and flagella tracking software

3D track reconstruction was performed in Matlab (The Mathworks) using an adapted version of the code from Taute et al.[17]. For automated analysis of the reorientation angle distribution, the script was successfully tested against simulated data with known reorientation angles (See SI). The helix parameter determination was performed automatically on tracks extending a duration of 0.4 s, with a mean-square-displacement of at least 10 µm^2^ and a mean opening angle between all consecutive velocity vectors of less than 60 ° to exclude strongly irregular tracks. The same exclusion parameters have been used for the reorientation angle analysis. Simultaneous dark-field cell and flagella tracking was performed using an in-house semi-automatic Matlab program. Tracking of the cell trajectory at 1640 fps without the flagella movement was performed using the TrackMate plugin of ImageJ.

### Hydrodynamic simulations

Each flagellum was modeled as a helical filament and a rotary motor. The helical filament was discretized with 20 beads with a discretization distance of 200 nm and bead diameter of 50 nm. Excluded volume interactions between all particles are considered using a truncated Lennard-Jones potential. Hydrodynamic interactions are taking into account using Stokesian dynamics simulation method having the translational anisotropic friction coefficients of *γ*_║_ = 1.6 ∙ 10^−3^ pNs/μm^2^ and *γ*_⊥_ = 2.8 10^−3^ pNs/μm^2^ and the rotational friction of *γ*_*r*_ = 1.26 10^−6^ pNs for the flagellum beads and *γ*_*bt*_ = 6πη*R*_*b*_ and 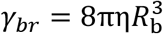 for the translational and rotational friction coefficients of the cell body. Irrespective of the high motor torque and isotropic bending and twisting rigidities mentioned in the main text, a stretching rigidity of 1000 pN, comparable to that of single flagella, could be chosen. A Rotne-Prager matrix was used for calculating the cross-mobilities and cross-hydrodynamics [27]. The swimming dynamics of the model cell at low Reynold number was calculated by solving the translational and rotational Stokes equations of motion for the cell body, flagellar beads and the bonds between them. A second-order Runge-Kutta algorithm [28, 29] and simulation time-steps of 10^−7^ s were used to solve the equations of motion numerically.

## Acknowledgements

The research leading to these results was supported by the Max Planck Society and by Deutsche Forschungsgemeinschaft (DFG) within the priority program on microswimmers (grants No. KL 818/2-2 and FA 835/7-2 to S.K. and D.F.). Further, S.M. was supported by Deutscher Akademischer Austauschdienst, DAAD (grant no. 57314018) as well as Deutsche Forschungsgemeinschaft (DFG) through SFB 937. A.C. is funded by the IMPRS on Multiscale Biosystems. C.T.L acknowledges support by the French National Research Agency (ANR Tremplin-ERC: ANR-16-TERC-0025-01). The authors would further like to thank L. Alvarez, R. Pascal and J. Jikeli for measurement trainings, C. Oschatz for medium preparation, A. Pohl for electron microscopy and C. Pilz for laboratory assistance.

## Supplementary Information

### Relation between helical and effective travel path

The overall traveled path l on a helix during one period can be parameterized using radius and pitch by 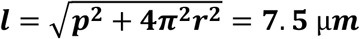. While effectively traveling 4 µm in 72 ms, the cell moves 7.5 µm on the large helix. While the contribution of the small helix to the actual swimming path provides another factor of 1.1, the ratio between instantaneous speed and effective speed is near 2.

### Discussion of pitch differences between simulation and observation

While the pitch is off by a factor of 2, the diameter and period time of helical trajectories are in agreement with experiment. The simulations show that the features of helical trajectories strongly depend on the flagellar opening angle (see Table S1) such that increasing the flagellar opening angle decreases the pitch and diameter of the large helix. Simulated effective velocities are smaller than the tracked velocities by factor of 2, since the pitch is decreased by the same factor while the period times are comparable. A detailed optimization of flagella opening angles and flagella length is likely to finally result in an exact agreement of measured and simulated pitches.

**Table S1.**
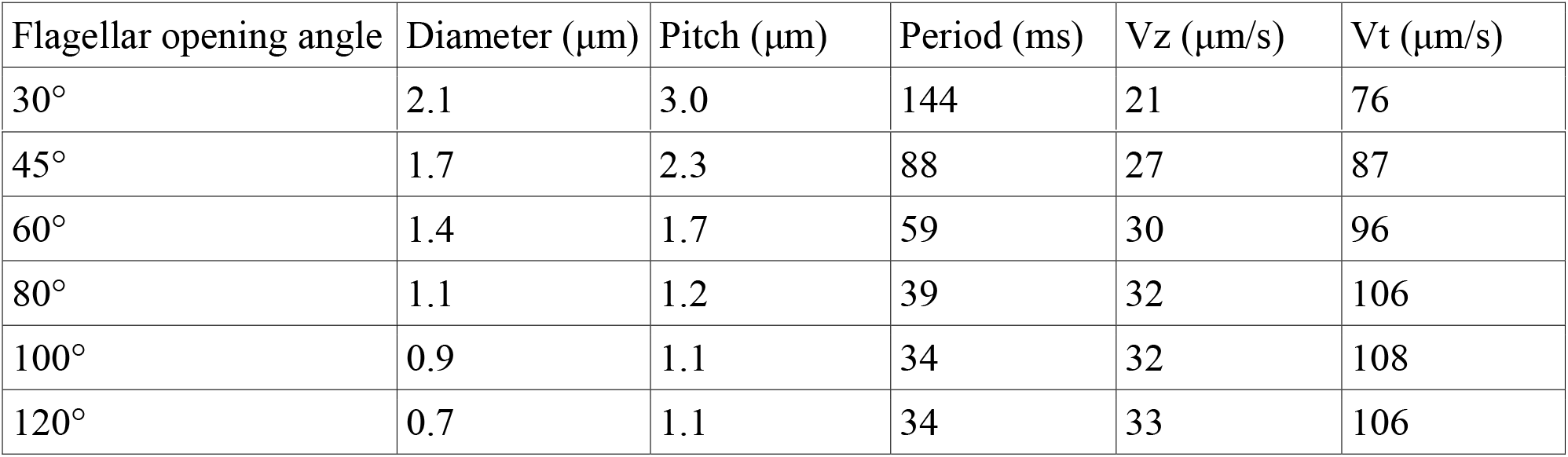
Swimming features of MC-1 cells using CCW&CW swimming mechanism for simulations with different flagellar opening angles. Given output parameters are the helix diameter, its pitch, the period time (period), the effective velocity (V_z_) and the instantaneous velocity (V_t_).

| Flagellar opening angle | Diameter ($\mu\text{m}$) | Pitch ($\mu\text{m}$) | Period (ms) | $V_z$ ($\mu\text{m/s}$) | $V_t$ ($\mu\text{m/s}$) |
| --- | --- | --- | --- | --- | --- |
| 30° | 2.1 | 3.0 | 144 | 21 | 76 |
| 45° | 1.7 | 2.3 | 88 | 27 | 87 |
| 60° | 1.4 | 1.7 | 59 | 30 | 96 |
| 80° | 1.1 | 1.2 | 39 | 32 | 106 |
| 100° | 0.9 | 1.1 | 34 | 32 | 108 |
| 120° | 0.7 | 1.1 | 34 | 33 | 106 |

### Flagella morphology

The individual bundles of the closed relative MO-1 [11] contain 7 individual flagella, which emerge from a hexagonal pattern on the cell envelope. Additionally, 24 gap-filling, presumably friction-reducing microfibrils were found.

**Figure S1:**
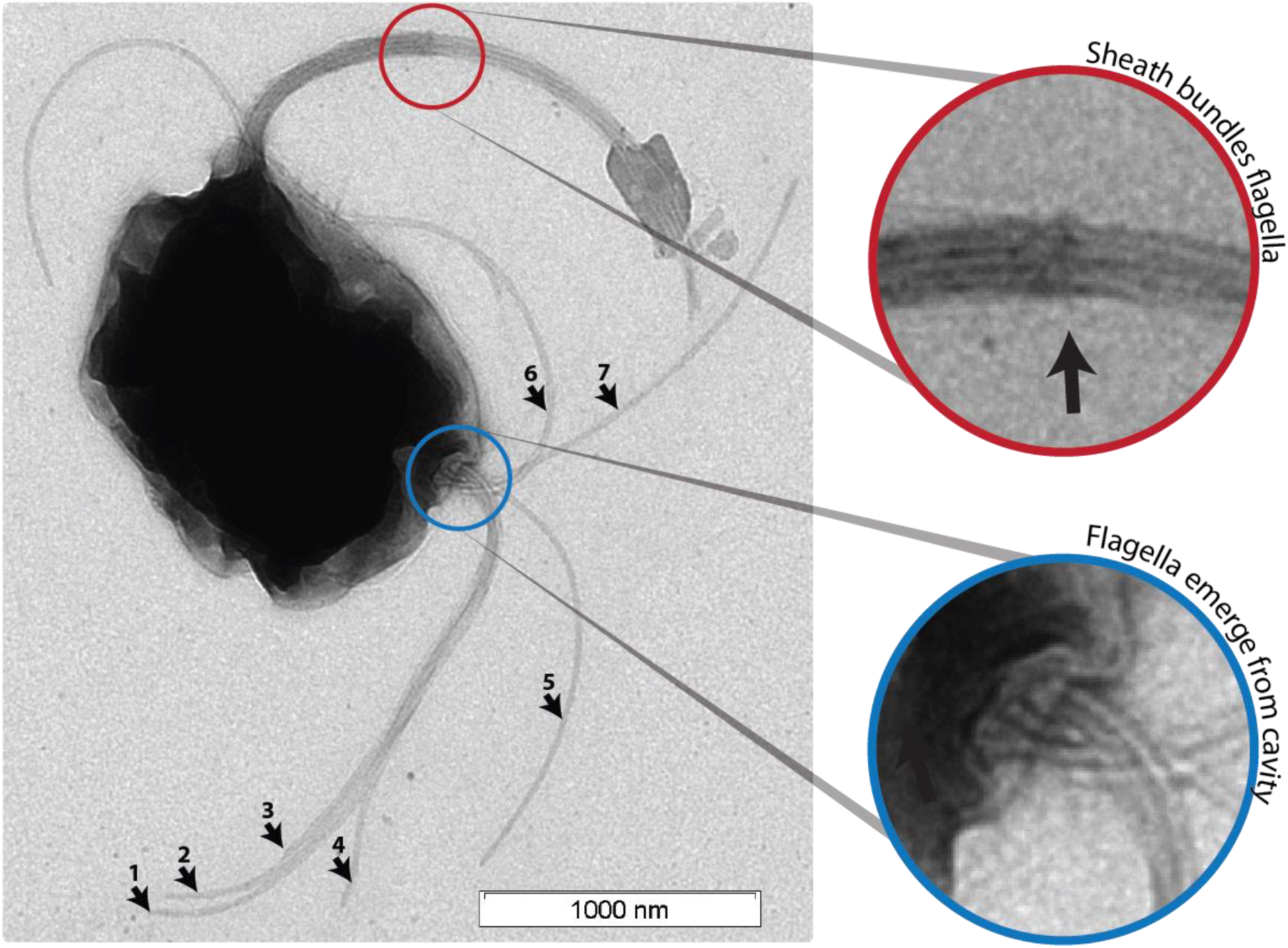
TEM images of an MC-1 cell stained with uranyl acetate. Two flagella bundles emerge from the MC-1 cell body. Each bundle features 7 flagella, wich emerge from a cavity on the cell surface. The individual flagella are bundeled by a sheath.

### 3D tracking validation

The algorithm for 3D track event detection was validated using Langevin-simulations of single active Brownian particles with defined mean reorientation angles, velocities, mean run times and mean run lengths, as quantified in Figure S2. The measurement evaluation is shown in Figure S3.

**Figure S2:**
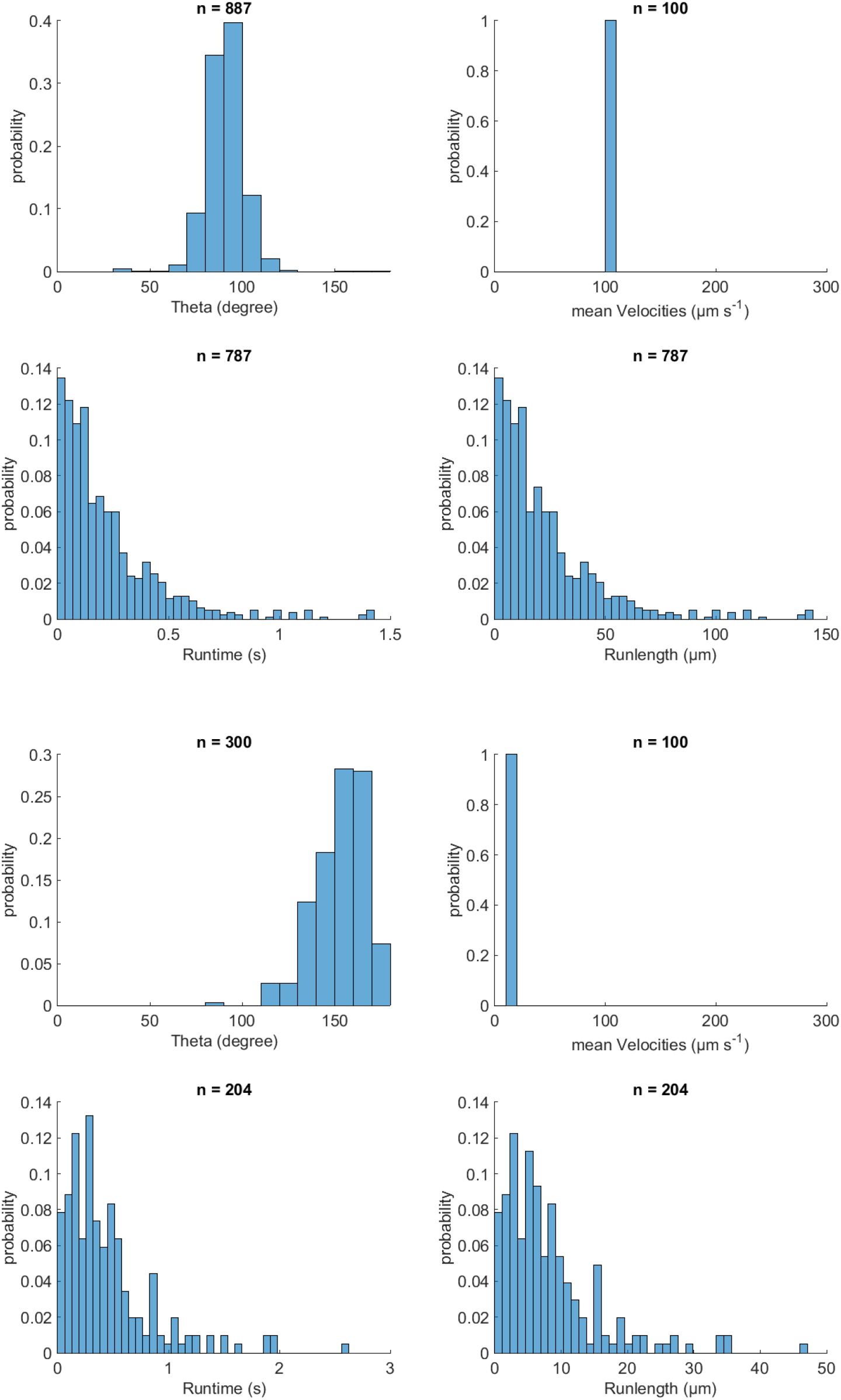
Validation of event identification script for 3D tracking on simulated tracks with known parameters. Top simulation parameters: Theta was 90 °, velocity was 100 µm/s, and runtime was 0.86 s. Bottom parameters: Theta was 170 °, velocity was 14 µm/s and runtime was 0.86 s. The mean time each reorientation lasted was 0.14 s for both cases.

**Figure S3:**
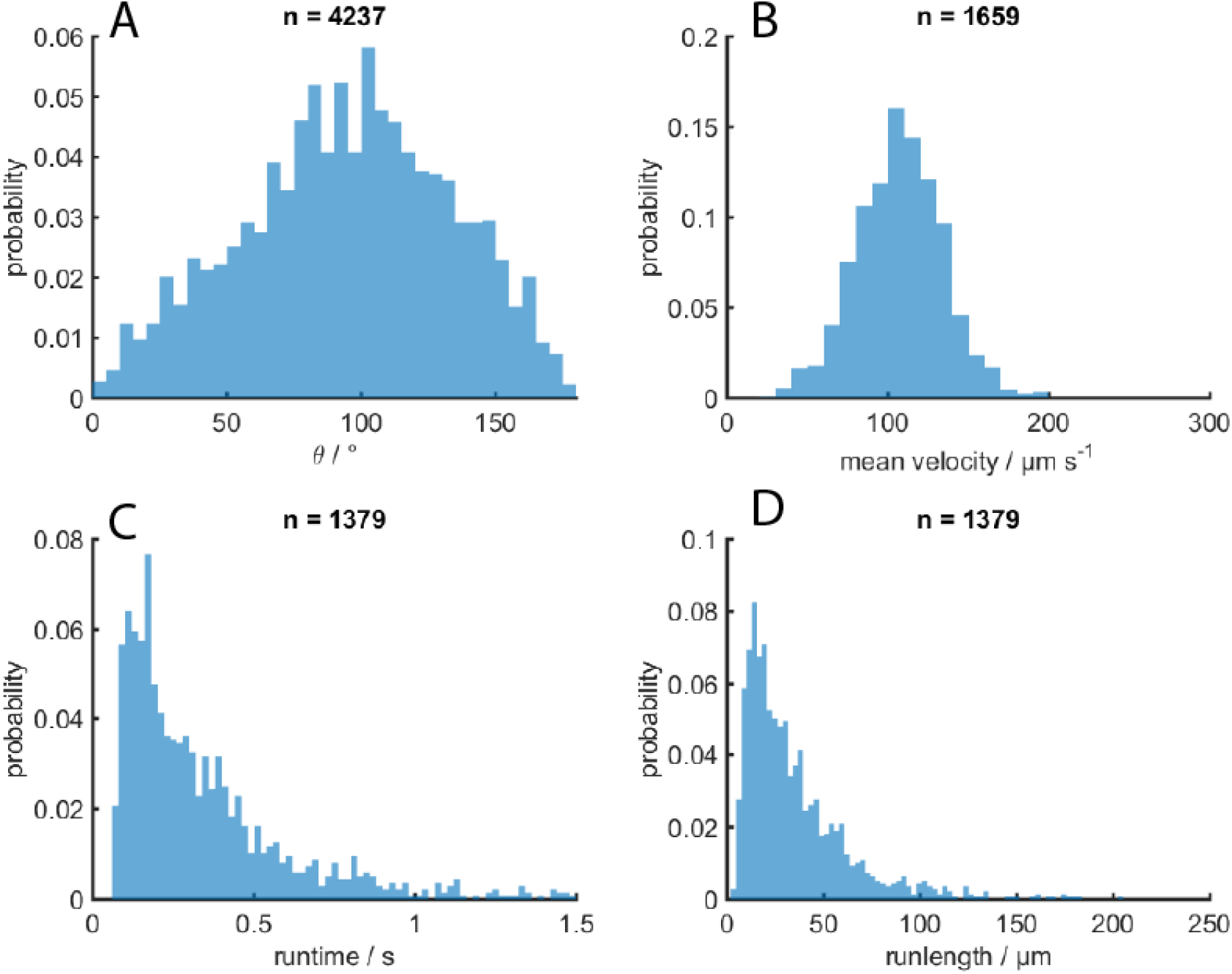
MC-1 swimming statistics. A) turning angle after a reorientation event, B) effective traveling speed, C) runtimes and D) runlengths in-between events.

**Figure S4:**
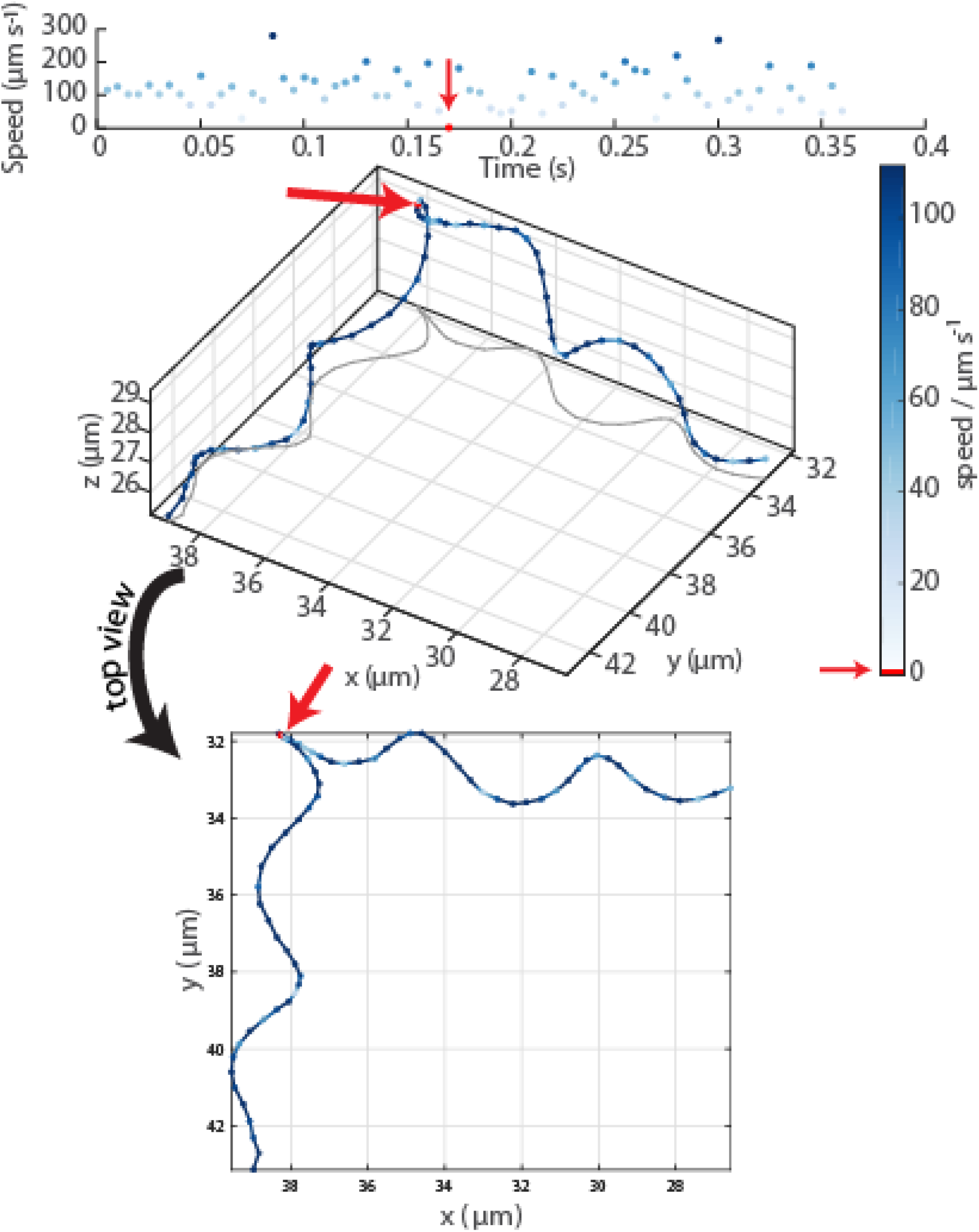
Oblique and top view of a typical event in an MC-1 swimming track. The radius, pitch and period time of the helix remain constant before and after the event, which is identified as a drop in velocity to 0 and which lasts 3.75 ms ± 1.25 ms.

### Test of different motor torque, magnetic moment direction and magnetic field configurations

**Figure S5.**
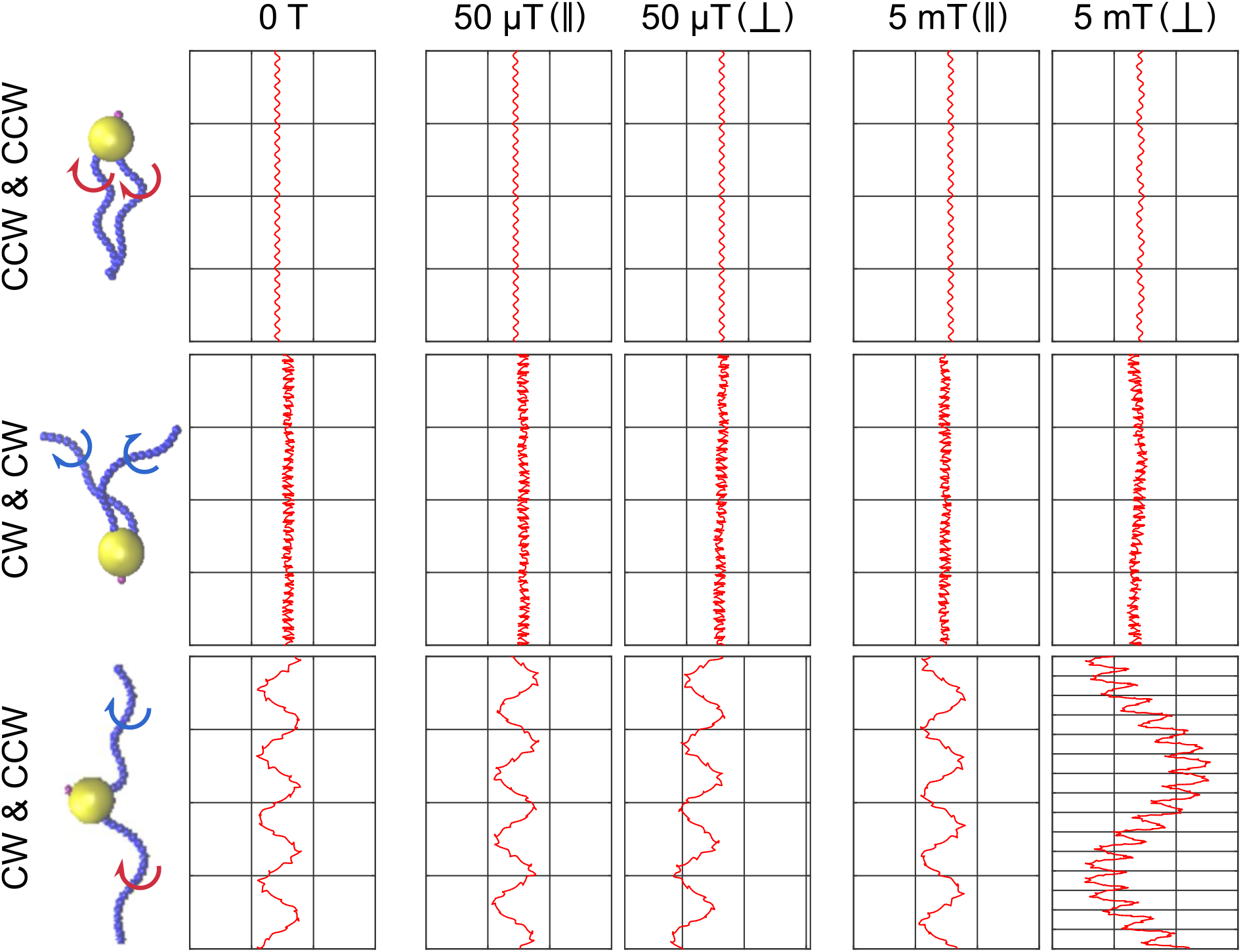
Simulations of different scenarios of MC-1 swimming. The tested flagella rotation patterns are shown together with the resulting cell trajectories for 5 different conditions: no magnetic field, weak magnetic field (50 μT) for a cell with magnetic moment parallel and perpendicular to the flagellar axes, and strong magnetic field (5 mT), again with magnetic moment parallel and perpendicular to the flagellar axes.. In the absence of a magnetic field, two pushing flagella produce small helices, two pulling flagella cause a strongly distorted movement pattern and only a cooperatively pushing and pulling flagella combination reproduces the experimentally observed double helical motion shown in Fig. 2A. A strong magnetic field results in the hyper-helical motion shown in Fig. 4D, if the magnetic moment is not parallel to the flagellar axes.

To check for the validity of simulation prediction, an experiment is setup in low and high magnetic fields, 50 *μT* and 3 mT, respectively. Over 1000 tracks are extracted and investigated for each magnetic field. For more certainty, swimming trajectories are investigated both in oxic and anoxic regions. In high magnetic field, over 80 trajectories can be detected illustrating the hyper-helix, while in Earth magnetic field only a few trajectories with semi-hyper-helix can be observed. A Typical MC-1 trajectories with the mentioned hyper-helix resulted from simulation and experiment are shown in Figure 4D. A measured diameter and pitch of the hyper-helix, *D*_exp_ ≃ 4.2*μm* and *P*_exp_ ≃ 30*μm*, are in good agreement with the simulated one, *D*_sim_ ≃ 3.9*μm* and *P*_sim_ ≃ 19.1*μm*.

## Video description

### Mov_SI01

Cell observation in a high-intensity dark-field video microscopy experiments at 1424 fps and 60× magnification with cell tracking.

### Mov_SI02

Cell observation in a high-intensity dark-field video microscopy experiments at 1424 fps and 60× magnification with cell tracking and tracking of white spots on the cell surface.

### Mov_SI03

Simulation of cell swimming with two pushing flagella.

### Mov_SI04

Simulation of cell swimming with one pulling and one pushing flagella.

### Mov_SI05

Example simulation of reorientation events.

